# Sox10 is required for systemic initiation of bone mineralization

**DOI:** 10.1101/2024.07.24.604990

**Authors:** Stefani Gjorcheska, Sandhya Paudel, Sarah McLeod, Louisa Snape, Karen Camargo Sosa, Cunming Duan, Robert Kelsh, Lindsey Barske

## Abstract

Heterozygous variants in the gene encoding the SOX10 transcription factor cause congenital syndromes affecting pigmentation, digestion, hearing, and neural function. Most of these symptoms are attributable to failed differentiation and loss of neural crest cells. Extensive research on mouse and zebrafish models has confirmed that Sox10 is essential for most non-skeletal crest derivatives, but seemingly dispensable for skeletal development. We challenge that concept here by revealing a novel requirement for Sox10 in skeletal mineralization. Neither neural crest- nor mesoderm-derived bones initiate mineralization on time in zebrafish *sox10* mutants, despite normal osteoblast differentiation and matrix production. We show that mutants are deficient in the ionocyte subpopulation tasked with taking up calcium from the environment through the Trpv6 epithelial calcium channel, leading to a severe calcium deficit that explains the lack of mineralization. As these ionocytes do not derive from a *sox10*+ lineage, we hypothesized that the primary defect instead resides in a separate organ that regulates ionocyte numbers or calcium uptake at a systemic level. Screening of the endocrine hormones known to regulate calcium homeostasis in adult vertebrates revealed significantly elevated levels of stanniocalcin (Stc1a), an anti-hypercalcemic hormone, in larval *sox10* mutants. Previous studies demonstrated that Stc1a inhibits calcium uptake in fish by repressing *trpv6* expression and blocking proliferation of Trpv6+ ionocytes. Our epistasis assays indicate that excess Stc1a is the proximate cause of the calcium deficit in *sox10* mutants. Lineage tracing shows that the pronephros-derived glands that synthesize Stc1a interact with *sox10*+ neural crest-derived cells, and that the latter are missing in mutants. We conclude that a subpopulation of Sox10+ neural crest non-cell-autonomously limit Stc1a production to allow the inaugural wave of calcium uptake necessary for the initiation of bone mineralization.

## Introduction

Sry-box transcription factor 10 (SOX10) is essential for pigmentation of the hair and skin, the ability to perceive sound and smell, and for digestive peristalsis. People with only one functional copy of the *SOX10* gene present pigment anomalies such as iris heterochromia and a white forelock, sensorineural hearing loss, deficient enteric innervation, anosmia, neurological abnormalities, neuropathy, and/or stalled puberty^1^. Cases range from mild to potentially lethal and are assigned to one of four congenital syndromes with overlapping clinical features: Waardenburg syndrome types 2E and 4C, Kallmann syndrome, or PCWH (Peripheral demyelinating neuropathy, Central dysmyelination, Waardenburg syndrome, and Hirschsprung disease)^1^. Besides the inner ear and central nervous system phenotypes, these symptoms are largely attributable to failed neural crest (NC) differentiation. This transient, migratory population of embryonic cells gives rise to pigment cells, sensory and enteric neurons and glia, the adrenal medulla, and bone, cartilage, and connective tissues of the facial skeleton^2^. All NC cells (NCCs) activate *SOX10* expression upon specification, prior to migration. The cranial subpopulation destined to give rise to the facial skeleton turn it off upon reaching their destination in the pharyngeal arches^3,4^. The remaining, non-skeletal NC populations retain Sox10 expression longer to activate programs for differentiation into pigment, glia, and sensory or enteric neurons, among other cell types^5^; in mutants, migration and differentiation stall, and the cells die^6^. Sox10 is also expressed in differentiating chondrocytes of both neural crest and mesodermal origin, but it is not critical there, as cartilage develops normally in zebrafish *sox10* mutants^7^. Accordingly, decades of research on heterozygous patients as well as homozygous mouse and zebrafish models has culminated in the notion that SOX10 is essential for non-skeletal neural crest derivatives but dispensable for formation of the skeleton^6–10^.

Bones mineralize by packing an organic collagenous extracellular matrix (ECM) with hydroxyapatite crystals of calcium and phosphate in a highly ordered manner^11^. Mature bone-forming osteoblasts secrete collagen I/X-rich ECM as well as enzymes (e.g. Alkaline phosphatase, Phospho1) and accessory glycoproteins (e.g. Osteopontin, Osteonectin) involved in synthesis and organization of the hydroxyapatite crystals^12–14^. Failed osteoblast maturation, disturbed matrix formation, and calcium-phosphate imbalances can disrupt ossification^15,16^.

Endocrine factors, particularly parathyroid hormone, vitamin D, and calcitonin, work in concert to maintain calcium and phosphate homeostasis in adults through actions on bone, intestine, and kidney^17–21^. Adult vertebrates obtain calcium and phosphate for all their cellular needs via dietary sources, environmental uptake, and renal reabsorption, as well as by breaking down bone^22^. However, how the initial wave of calcium and phosphate uptake in the developing embryo is regulated remains a gap in knowledge. Mammalian fetuses obtain minerals largely through the placenta^23^, while fish larvae take them from maternal yolk stores or directly out of the water through ionocytes in the skin and gills^24,25^. Indeed, zebrafish larvae obtain the necessary amount of phosphate through phospholipid metabolism in the yolk and do not require additional environmental phosphate uptake^26–28^. Conversely, calcium uptake from the environment is required for skeletal mineralization^29^. The major route of calcium ingress is the constitutively open epithelial calcium channel (ECaC) encoded by *trpv6* (Transient Receptor Potential channel family, Vanilloid subfamily member 6)^29–31^. Of the five major types of ionocytes in fish, only the Na^+^/H^+^-ATPase-rich (NaR) subpopulation expresses *trpv6*^25,30^. *Trpv6* expression is also highly enriched in both maternal and fetal cells of the mammalian placenta^32^. Because Trpv6 is constitutively open, regulation of calcium uptake occurs through modulating levels of *trpv6* transcription or the proliferation/quiescence of *trpv6*+ cells^30^. Whether the major endocrine hormones involved in calcium and phosphate homeostasis in adults also control the initiation of calcium uptake via Trpv6 for skeletal mineralization in the embryo remains largely unknown.

One factor known to not drive but rather limit calcium uptake in both embryonic and adult fish is an anti-hypercalcemic hormone called stanniocalcin^33^. Stanniocalcin (Stc1) is a glycoprotein secreted by a variety of tissues in mammals (e.g. kidney, intestine), where it is involved in local calcium homeostasis^34–38^. Stc1a was first isolated from the Corpuscles of Stannius (CS), intermediate mesoderm-derived endocrine organs unique to teleost fish^35,39,40^. Surgical removal of the CS or mutation of the *stc1a* gene causes severe hypercalcemia, kidney stone formation, and an increase in NaR cell numbers in fishes^35,41,42^. Conversely, exposure to high environmental calcium increases *stc1a* mRNA levels and serum Stc1a content, in turn leading to decreased calcium uptake^33^. Stc1a’s anti-hypercalcemic activity involves inhibition of both *trpv6* expression and NaR cell proliferation, working through a Pappaa-Igfbp5a-Igf-Igfr cascade that impacts PI3K, mTor, and Akt signaling^42–46^. In normal or low calcium, active Pappaa cleaves the Igf-binding protein Igfbp5a, releasing Igf ligands to activate downstream signaling and NaR cell proliferation. In conditions of high environmental calcium, Stc1a inhibits Pappaa’s protease activity, keeping NaR cells quiescent^42–46^.

In this study, we present a previously undescribed systemic requirement for Sox10 in the initiation of skeletal mineralization in fish. We provide evidence of a striking Stc1a increase in *sox10* mutants that severely reduces *trpv6*+ ionocyte number and whole-body calcium content. We find *sox10*+ neural crest-derived cells interacting with the Corpuscles of Stannius in control but not mutant fish, indicating that they may serve to moderate *stc1a* levels in the embryo, allowing the massive wave of calcium uptake required to initiate bone mineralization

## Results

### Delayed onset of skeletal mineralization in zebrafish *sox10* mutants

Though Sox10 expression is activated in all NCCs upon specification^5,6,47,48^, it is quickly downregulated in the subset of cranial crest that will go on to form the facial skeleton^4^. These skeletal progenitor cells were presumed to not require Sox10 function, as our early studies noted no defects in the Alcian blue-labeled cartilages of the zebrafish *sox10* mutant facial skeleton^7^. However, while recently performing a routine bone stain in *sox10* mutants, we unexpectedly noticed a striking absence of mineralization when the skeleton is first differentiating at 3-4 days post fertilization (dpf) (Fig. 1A). At these stages, calcium deposits in newly mineralizing bones are readily apparent by Alizarin red staining in sibling controls. Weak staining appears in mutants by 5 dpf and increases until larval lethality around 8 dpf, but never attains control levels. Mutants are not edemic or developmentally delayed, ruling out these common explanations for poor mineralization. The phenotype is also indiscriminate of ossification type (endochondral, intramembranous, and even odontogenic) and bone developmental origin in the mesoderm versus neural crest (Fig. 1B).

**Figure 1.**
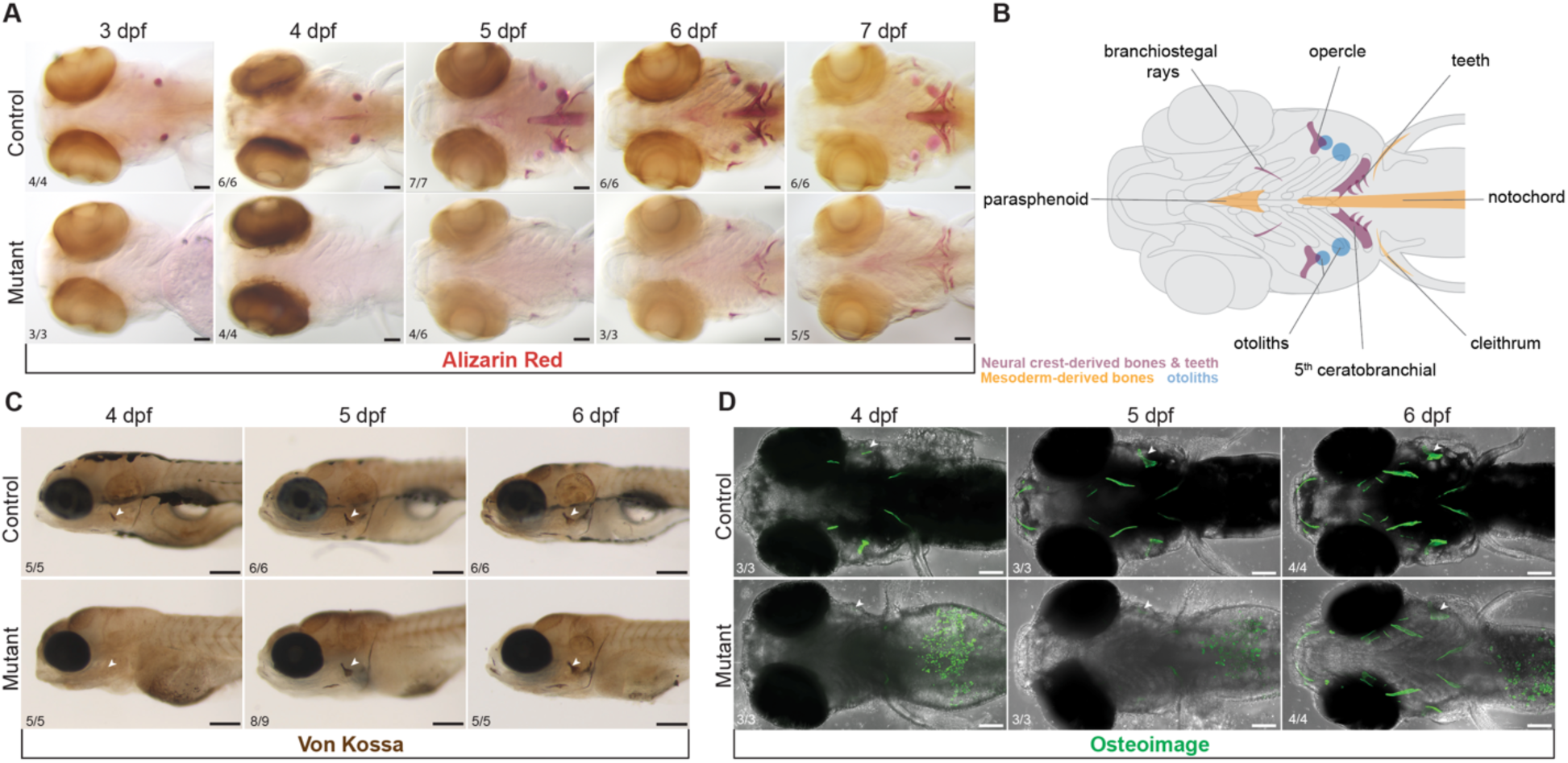
Mineralization deficit in zebrafish *sox10* mutants. **(A)** A major delay in initiation of bone mineralization in *sox10* mutants between 3 and 7 dpf is revealed by Alizarin red staining. Some mineralization is present by 5 dpf but never achieves control levels before lethality at 8 dpf. Scale bar: 100 µm. Schematic representation of the affected mineralized structures and their embryonic origins. **(C, D)** Von Kossa (C, scale bar: 200 µm), and Osteoimage (D, scale bar: 100 µm) staining show absent calcium deposition and hydroxyapatite formation in *sox10* mutants at 4 dpf and gradual recovery starting at 5 dpf. Arrowheads pointing at the opercle (op). Numbers in all panels indicate the proportion of larvae of that genotype with the presented phenotype.

As deficient mineralization has not been reported in any of the many existing mouse, fish, or frog Sox10 loss-of-function models, we questioned whether it could be a neomorphism specific to our *sox10^ci30^*^20^ allele^49^. *ci3020* is a 1495-bp deletion that removes part of the 5’UTR and the first coding exon, encoding the homodimerization domain and part of the DNA-binding high mobility group (HMG) domain (Fig. S1A). Some transcription still occurs from the deletion allele^49^, and the first in-frame methionine downstream of the deletion could conceivably produce an N-terminally truncated protein lacking the HMG box but retaining the transactivation domain^1^. *sox10^ci30^*^20^*^/ci3020^* embryos otherwise present the classic *colourless* phenotypes associated with *sox10* loss-of-function (Fig. S1B-C)^50^, lacking melanocytes and xanthophores, with malformed otic vesicles and otoliths but normal facial cartilages. To test whether deficient mineralization is specific to the *ci3020* allele, we performed Alizarin red staining on homozygotes for the *m618* (L142Q) missense allele first reported in 1996^51^. The same near-absence of staining was observed between 4 and 6 dpf (Fig. S1D), demonstrating that this phenotype is a general consequence of loss of *sox10* function, at least in zebrafish. We further validated the Alizarin red results in *ci3020* mutants (hereafter *sox10* mutants) using Von Kossa and Calcein stains (Fig. 1C, S1C’), which both label calcium deposits,^52–55^ as well as Osteoimage (Fig. 1D), a stain that specifically detects hydroxyapatite^56^. These stains confirmed that mineralization gradually initiates around 5 dpf, first apparent by Von Kossa staining (Fig. 1C). Supporting that the recovery is incomplete, fluorescent Calcein staining in older 7 dpf larvae revealed a lack of endochondral bone collars around the mutant hyomandibula and ceratohyal cartilages (Fig. S1C’).

Osteoblasts are essential for mineralization, but do not themselves express Sox10 (Fig. 2A-B) ^57^. The subset of osteoblasts derived from the cranial neural crest did transiently express *sox10* and accordingly express the *SOX10:*Cre neural crest lineage label (by a human neural crest-specific *SOX10* promoter)^5,48^, but those derived from mesoderm never pass through a *sox10*+ state. We therefore presumed that the broad mineralization deficit would not be cell-autonomous to osteoblasts, though it was still possible that their differentiation could be impacted by extrinsic factors. Osteoblasts are evident as early as 3 dpf at the site of the future opercle (op) bone^58^. To evaluate mutant osteoblasts, we used established transgenic markers *RUNX2*:mCherry^59^, *sp7*:EGFP^60^ and *osc*:EGFP^29^, which are respectively activated in osteoprogenitors and early and maturing osteoblasts. Live imaging of the op bone in *sox10* mutants and sibling controls from 3 to 7 dpf revealed seemingly normal patterns for each marker (Fig. 2C-C’’). Visualization of *sp7:*GFP in combination with live Alizarin red staining confirmed that individual elements are growing similarly between mutants and controls (Fig. 2D). Colorimetric *in situs* for the major bone ECM component *col10a1a* also revealed normal expression in mutants (Fig. 2E). These findings suggest that mutant osteoblasts are still differentiating and making collagenous matrix despite not being able to mineralize it.

**Figure 2.**
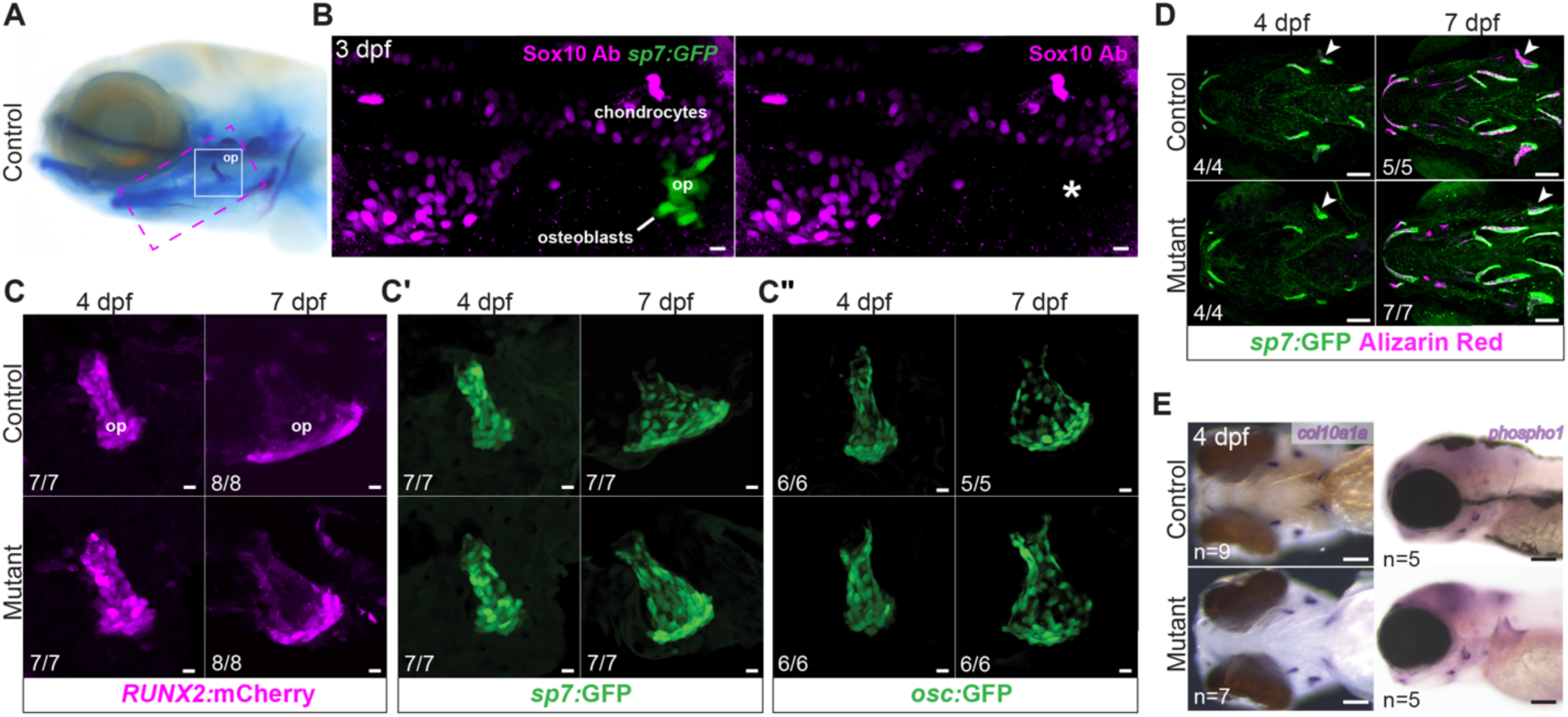
Normal patterns of growth and differentiation in *sox10* mutant osteoblasts. **(A)** Reference image of a larva stained with Alcian blue and Alizarin red, with locations of skeletal elements shown in B (magenta dashed line) and C (white line) highlighted. op, opercle. **(B)** Immunostaining with an anti-Sox10 antibody reveals strong expression in chondrocytes but a lack of Sox10 protein (asterisk) in mineralizing osteoblasts (*sp7*:GFP+) forming the op bone at 3 dpf. Scale bar: 10 µm. **(C)** Normal growth of *sox10* mutant op (arrowhead) as well as other bones despite minimal calcium accumulation, revealed by live imaging of Alizarin red-stained *sp7*:GFP+ embryos at 4 and 7 dpf. Scale bar: 100 µm. **(D-D’’)** Sequential live imaging shows normal patterns of *RUNX2*:mCherry, *sp7*:GFP and *osc*:GFP transgene expression in mutant osteoblasts of the op at 4 and 7 dpf. Scale bar: 10 µm. **(E)** Colorimetric *in situ* hybridizations for *col10a1a* and *phospho1*, encoding key bone matrix components, revealed no overt abnormalities in *sox10* mutants at 4 dpf. Scale bar: 100 µm.

To determine whether the mineralization machinery is intact in *sox10* mutant osteoblasts, we performed *in situs* and/or semi-quantitative rt-PCR for *phospho1*, *spp1* (Osteopontin), *sparc* (Osteonectin), *alpl* (Alkaline phosphatase), *enpp1*, *entpd5*, *phex*, *fgf23*, *runx2a*, and *runx2b* (Fig. 2E, S2)). These genes encode for secreted proteins and enzymes associated with matrix formation, phosphate and calcium regulation, and hydroxyapatite synthesis, in addition to the Runx2 transcription factors required for osteoblast specification. In rt-PCRs performed on cDNA made from pooled 4-dpf embryos, we detected mild increases in *alpl* and *entpd5* in the mutants (p<0.05, unpaired t-tests; Fig. S2A). We also observed slight decreases in *spp1*, *phospho1*, *enpp1*, and *fgf23* in the mutants (p<0.05, unpaired t-tests; Fig. S2A), a pattern opposite than observed in the zebrafish *enpp1* mutant, which shows increased mineralization^61^. There was no change in *sparc* or *phex*^61^ (p>0.0.5, unpaired t-tests) in mutant compared to wild-type embryos (Fig. S2A). *In situ* hybridizations revealed unchanged *runx2a* and *runx2b* expression (Fig. S2B), aligning with the live-imaging *RUNX2:*mCherry experiment (Fig. 2D). *spp1* was strikingly reduced, consistent with the rt-PCR result (Fig. S2A-B). However, inconsistent with the rt-PCR results, *phospho1* expression in forming bones appeared largely unchanged in mutant heads (Fig. 2E), while *sparc* appeared reduced (Fig. S2B). Discrepancies may be due to altered expression in other tissues not captured by the *in situs.* These findings nonetheless demonstrate for the first time that multiple factors linked with mineralization anomalies in animal models and human patients^62–65^ are dysregulated in *sox10* mutants.

### *sox10* mutants are calcium-deficient

Disrupted mineral homeostasis caused by mutations in the phosphate regulators *enpp1* and *entpd5* impacts the expression of many other mineralization-regulating factors, including many of the genes we assayed^61,62^. To test whether the observed dysregulation in our mutant could also be a consequence of a systemic mineral imbalance, we measured calcium and phosphate levels in our mutants. As it is not possible to directly measure serum mineral contents in larval fish, we used a colorimetric assay (Fig. 3B) on pooled whole-body samples between 36 and 168 hpf, following standard practice in the field^33,42,66^. In wild-type zebrafish, Ca^2+^ content begins to increase around 3 dpf as the first bones mineralize and continues to rise with age (Fig. 3A)^67^. By contrast, *sox10* mutants had lower Ca^2+^ content compared with controls starting at 3 dpf (p=0.03, unpaired t-test; Fig. 3A-B). Consistent with the bone staining in mutants first appearing around 5 dpf (Fig. 1A), we found that mutant Ca^2+^ levels at 5 dpf were approximately equivalent to control levels at 3 dpf (0.01 mg/embryo), suggesting this may be the minimal Ca^2+^ threshold required to initiate mineralization. To further investigate this possibility, we raised wild-type embryos in medium completely devoid of Ca^2+^ and found that mineralization was absent everywhere except the otoliths inside the otic vesicles (Fig. S3A’). These are made of calcium carbonate rather than hydroxyapatite^68,69^ and also still form in *trpv6* mutants that cannot take up external calcium^29,30^. The Ca^2+^ content of these wild-type fish raised in 0 mM Ca^2+^ was approximately the same as that of mutants raised in 1 mM Ca^2+^ (Fig. S3A), supporting that this low level is below the threshold needed for bone mineralization. On the other hand, phosphate levels were seemingly unaffected in mutants between 36 and 168 hpf (Fig. S3B), suggesting that lack of calcium is the major cause of the delayed and deficient hydroxyapatite formation (Fig. 1D).

**Figure 3.**
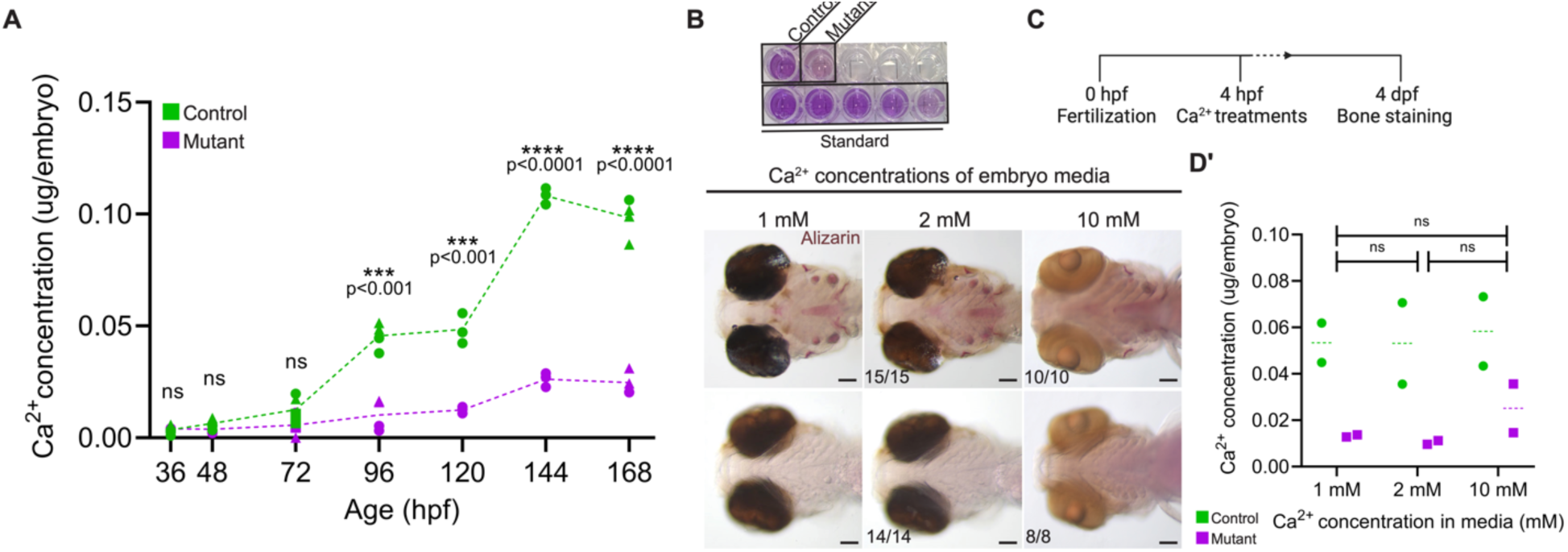
*sox10* mutants have a severe whole-body calcium deficit. **(A-B)** Colorimetric calcium assay reveals significantly lower levels of Ca^2+^ in *sox10* mutants after mineralization is initiated at 3 dpf. Each data point represents a pool of 10-15 embryos. Different shapes represent biological replicates assayed on different days (unpaired t-tests: 36 hpf: p=0.580, df=8; 48 hpf: p=0.083, df=8; 72 hpf: p=0.091, df=7; 96 hpf: p=0.0002, df=6; 120 hpf: p=0.0008, df=4; 144 hpf: p=0.000008, df=4; 168 hpf: p=0.000005, df=6). B is an example of the colorimetric assay, showing a clear reduction in mutants. **(C)** Schematic representation of the Ca^2+^ treatment protocol. **(D-D’)** Increasing ambient Ca^2+^ levels to 2 or 10 mM does not rescue the mineralization deficit (D; scale bar: 100 µm) or Ca^2+^ content (D’) (unpaired t-tests: 1 vs. 2 mM: p=0.963, df=2; 1 vs. 10 mM: p=0.778, df=2; 2 vs. 10 mM: p=0.748, df=2). Ratios reflect the number of imaged larvae of that genotype with the presented phenotype. In D’, bars indicate the median; significance determined by unpaired t-test. Scale bar: 100 µm.

Other zebrafish mutants with poor mineralization but seemingly normal osteoblasts, e.g., *msp* and *trpv6*, can be rescued by simply increasing the concentration of Ca^2+^ in the media^29,70^. We tested whether this would also improve our phenotype using Ca^2+^ concentrations two- and ten-fold higher than our standard embryo media (2 and 10 mM versus 1 mM, respectively, following^62,70^ (Fig. 3C-D)). However, Alizarin red staining at 4 dpf revealed no increase in mineralization in mutants reared in either high-Ca^2+^ medium (Fig. 3D). We then quantified their Ca^2+^ contents at 4 dpf to specifically assess the calcium deficit, finding that mutants raised in the highest-Ca^2+^ environment did show a non-significant increase in Ca^2+^ content, but they remained at a severe deficit relative to controls (Fig. 3D’). Lowering or increasing the phosphate concentration likewise had no impact on mineralization in mutants (Fig. S3C). The mineralization delay in the *sox10* mutants may thus have a more complex etiology than other mutant lines with similar phenotypes.

Calcium is taken up from the environment in fish larvae through Trpv6 channels present on the surface of specialized NaR ionocytes in the skin^71^. NaR cells also uniquely express *igfbp5a*^44,72^ and comprise a subset of ionocytes expressing Na^+^/K^+^ ATPase^25^. Immunostaining for Na^+^/K^+^ ATPase combined with the *SOX10:*Cre lineage label (driven by a human neural crest-specific enhancer^5,48^) in otherwise wild-type fish confirmed that these skin ionocytes do not derive from neural crest (Fig. 4A), in line with previous work tracing them to the ectoderm^73^. We questioned whether the persistently low calcium content of *sox10* mutants could be due to a deficiency of *trpv6* expression or total NaR ionocytes. Though rt-PCR revealed no overt change in whole-body *trpv6* levels (Fig. 4B), we did detect significant decreases in the numbers of *trpv6+* and *igfbp5a*+ cells at 4 dpf, with mild recovery by 7 dpf (Fig. 4C-D’). These patterns support reduced NaR number (rather than *trpv6* transcription) as the cause of the systemic calcium deficit and the associated lack of bone mineralization. Published scRNAseq data confirm that differentiating NaR cells contain no *sox10* transcripts^74^. The NaR deficit in *sox10* mutants therefore cannot be explained by a simple cell-autonomous requirement for Sox10.

**Figure 4.**
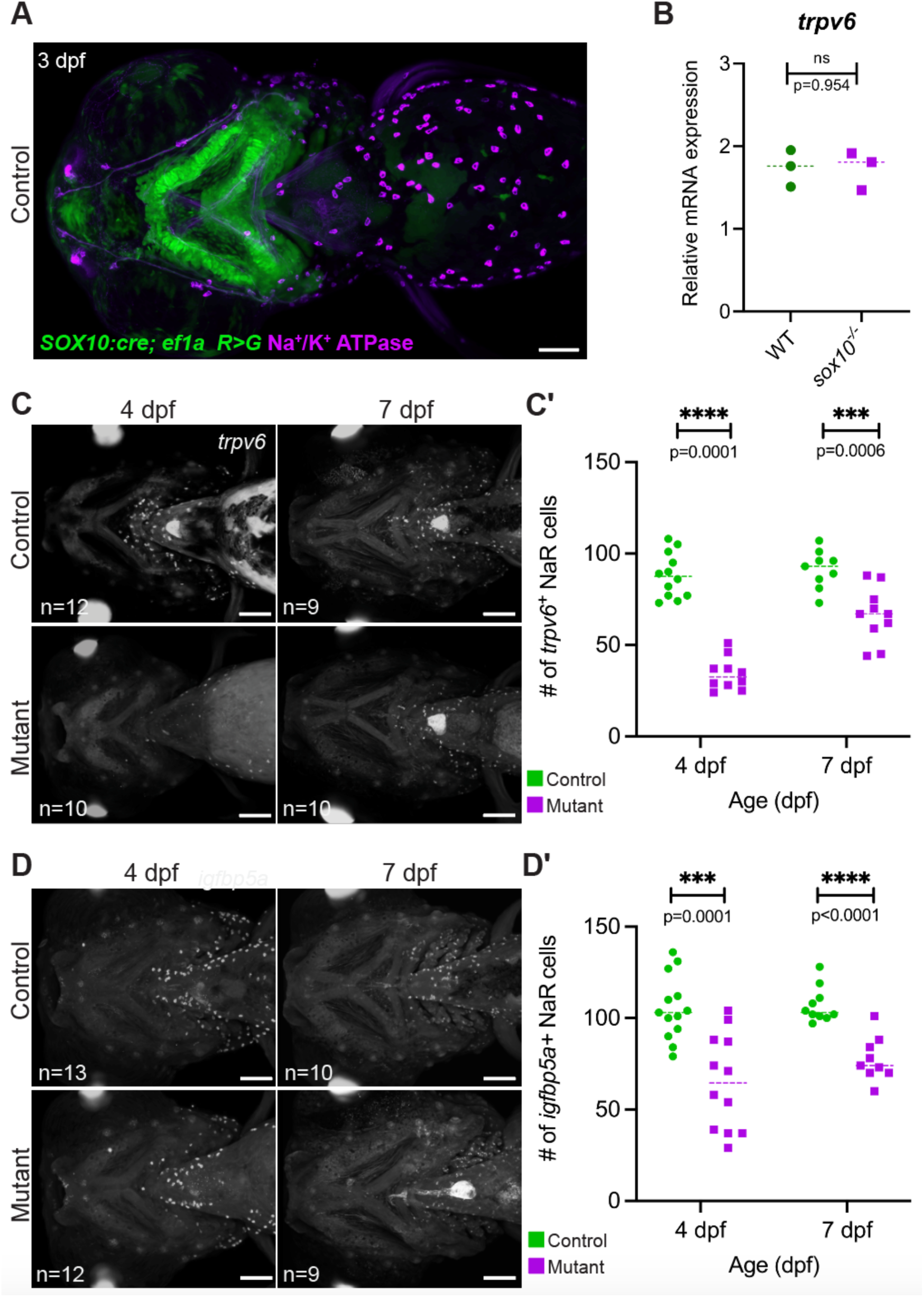
Reduction in *trpv6*+ NaR cell number in the *sox10* mutants. **(A)** Immunostaining of a 3 dpf *SOX10:Cre; ef1a: DsRed>GFP* larva with an antibody against the Na+/K+ ATPase pump confirms that NaR ionocytes do not derive from neural crest. Scale bar: 100 µm. **(B)** rt-PCR demonstrates that *trpv6* transcription is not overtly altered at the whole-body level at 4 dpf. Each point represents a pool of 10-15 embryos (unpaired t-test: p=0.954, df=4). **(C-D’)** Fluorescent *in situ* hybridizations for *igfbp5a* and *trpv6* (C,D) both demonstrate a striking and significant reduction in the number of NaR cells in mutants at 4 dpf (quantified in C’, D’), with partial recovery by 7 dpf (unpaired t-tests; *trpv6*: 4 dpf: p<0.000001, df=20; 7 dpf: p=0.0006, df=17; *igfbp5a:* 4 dpf: p=0.0001, df=23; 7 dpf: p<0.0001, df=17). Scale bar: 100 µm.

NaR cell numbers fluctuate depending on the amount of calcium in the environment, with low Ca^2+^ stimulating their proliferation and thereby increasing Ca^2+^ uptake, versus minimal proliferation and uptake under high Ca^2+^ ^24,75^. These fluctuations still occur in *sox10* mutants (Fig. S4A), indicating that they are still capable of responding to environmental conditions. However, the increase in NaR cells observed in mutants raised at low Ca^2+^ is dampened relative to controls, apparently insufficient to raise total Ca^2+^ content (Fig. S3A) or permit robust skeletal mineralization (Fig. S3A’).

### Endocrine suppression of NaR ionocyte expansion in *sox10* mutants

The fact that the number of *trpv6*+ NaR cells remains so low in *sox10* mutants despite their clear need for calcium struck us as paradoxical. We reasoned that mutants might be lacking a factor needed to stimulate NaR proliferation, or, conversely, have too much of a different factor that blocks their increase. In an rt-PCR screen of candidate endocrine factors, we identified stanniocalcin-1a (*stc1a*) as being 3-fold upregulated in *sox10* mutants at 4 dpf (Fig. 5A). Stc1a is an anti-hypercalcemic hormone triggered by high environmental calcium through activation of the Calcium-Sensing Receptor (CaSR)^33,76–78^. Stc1a reduces calcium uptake to maintain physiologically safe levels by inhibiting proliferation of NaR cells and suppressing *trpv6* expression^42,45^. The dominant sources of Stc1a in fish larvae are the Corpuscles of Stannius, teleost-specific glands that bud off the distal pronephros by 50 hpf and are positioned to either side of the posterior cardinal vein with their own vascular supply by 3 dpf (Fig. 6A-B)^40,79,80^. *stc1a* expression is detectable prior to completion of CS extrusion^40^ and is thus potentially involved in maintaining calcium balance as early as 24 hpf^40^.

**Figure 5.**
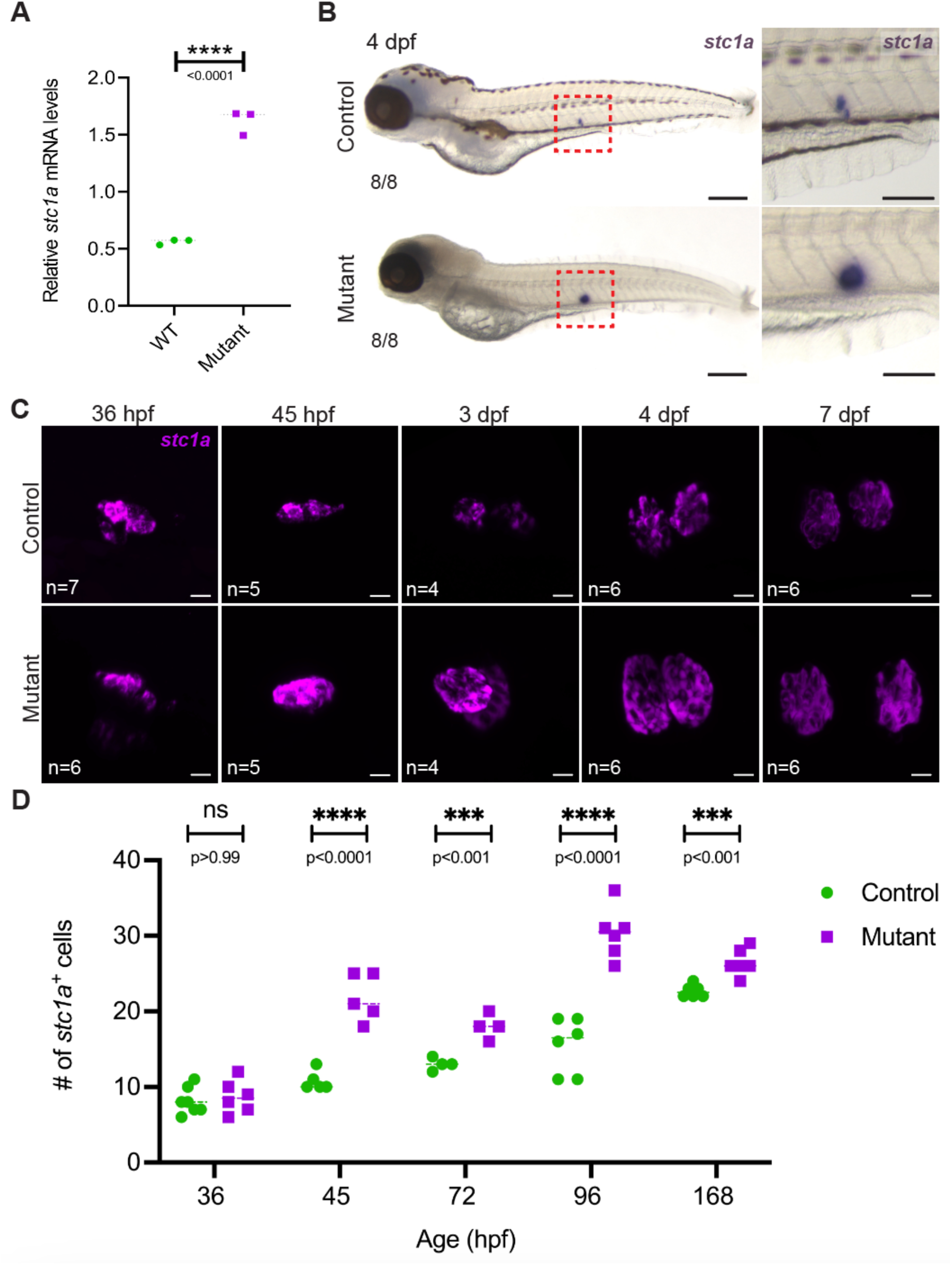
Upregulation of anti-hypercalcemic hormone *stc1a* in *sox10* mutants. (A-B) Both semi-quantitative rt-PCR (A) and *in situ* hybridization (B) detect a robust upregulation of *stc1a* mRNA in *sox10* mutants at 4 dpf (unpaired t-test in A, p<0.0001, df=4). Scale bars: 200 µm in B and 100 µm in inset. (C-D) The increase in *stc1a* transcript levels is due at least in part to an increase in the number of *stc1a+* cells in *sox10* mutant Corpuscles, first detected at 45 hpf and resolving at 7 dpf (unpaired t-tests; 36 hpf: p=0.640, df=11; for 45 hpf: p=0.00009, for df=8; for 72 hpf: p=0.002, df=6; for 96 hpf: p=0.00003, df=10; for 168 hpf: p=0.0007, df=10. Scale bars in C: 10 µm.

**Figure 6.**
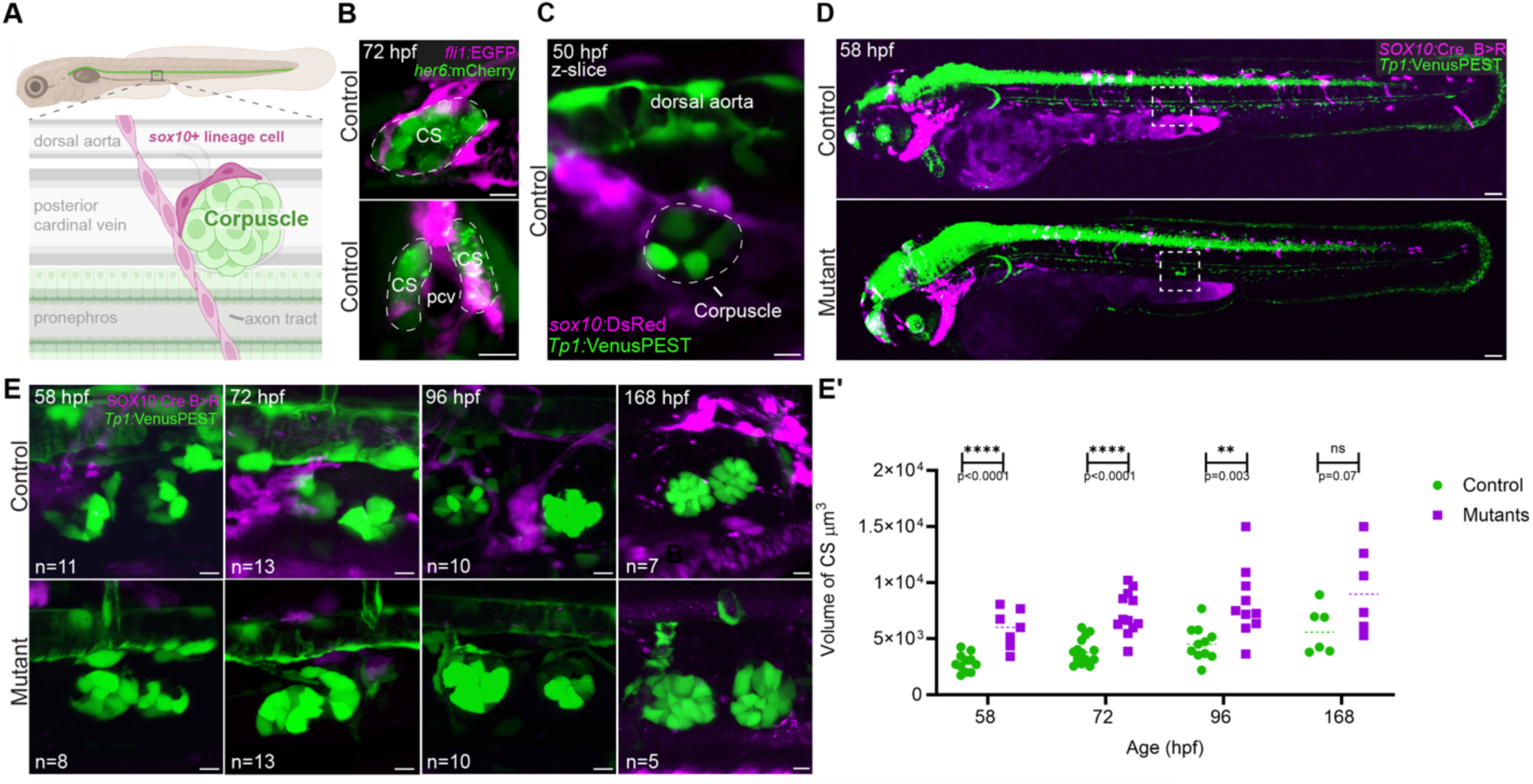
*sox10*+ crest-derived cells surround the Corpuscles of Stannius and are missing in *sox10* mutants. **(A)** Schematic representation of the Corpuscles’ position and surrounding environs. The glands are positioned ventral to the dorsal aorta (da), dorsal to the pronephros, and flanking the posterior cardinal vein (pcv). *sox10+* lineage cells branch off neighboring axon tracts (lined with sox10+ Schwann cells and Schwann cell precursors) to contact the Corpuscles. **(B)** Control 72 hpf larva doubly transgenic for *fli1:*EGFP (magenta, labeling vasculature) and *her6:*mCherry (green, labeling the Corpuscles). Top image is a single optical section from the lateral perspective showing endothelial cells wrapping around the CS. Bottom image is a maximum intensity projection rotated orthogonally to show the close interaction of the CS with the pcv. **(C)** Single optical section showing that *sox10:*DsRed+ cells are also in close contact with the *Tp1*:VenusPEST+ Corpuscles by 50 hpf. **(D-E)** Whole-mount live imaging of *SOX10:Cre B>R; Tp1*:VenusPEST fish shows *sox10*+ lineage cells surrounding the CS at earlier stages (58 and 72 hpf), nearby at 96 and 168 hpf, and absent in the *sox10* mutants. The entire fish is shown in D to highlight the deficiency of trunk NCCs (magenta). **(E’)** Volume measurements of control and mutant *Tp1:*VenusPEST+ CS between 58 and 168 hpf. Only the right CS, closest to the lens, was measured (unpaired t-tests; 58 hpf: p=0.0001, df=16; 72 hpf: p=0.00001, df=24; 96 hpf: p=0.003, df=19; 168 hpf: p=0.068, df=10). Scale bars: B and D: 10 µm, C: 100 µm.

Aberrantly elevated *stc1a* expression in *sox10* mutants might thus explain their reduced number of NaR cells and calcium uptake. In situ analyses showed that the robust increase first becomes apparent after completion of CS extrusion (after 36 hpf; Fig. 5C), is most obvious at 4 dpf (Fig. 5B-C), then begins to level out by 7 dpf (Fig. 5C), when both *trpv6+* NaR cell numbers and mineralization are partially recovering. The *stc1a* increase is due at least in part to higher numbers of *stc1a+* cells in the mutant CS between 45 hpf and 4 dpf (p<0.001, unpaired t-test; Fig. 5D). Interestingly, in low-Ca^2+^ medium, *stc1a* expression is undetectable in controls but merely reduced in mutants (Fig. S4B), possibly explaining why mutants still have fewer NaR cells and less calcium uptake than their siblings under these conditions (Fig. S3A, S4A).

The *stc1a*-expressing Corpuscles are derived from intermediate mesoderm^40,81^ and never pass through a *sox10+* state, so their dysfunction in *sox10* mutants must also be indirect. We looked for *sox10* lineage+ cells in or surrounding the glands, predicting that they may be aberrant or missing in mutants. We tracked neural crest using *SOX10*:Cre^5,48^ in combination with the *actb2:*BFP*>*DsRed Cre reporter^82^ and all recently *sox10*-expressing cells using *sox10*:DsRed (driven by the 4.9-kb zebrafish *sox10* promoter^83^). All traces were performed in combination with the *Tp1:*VenusPEST Notch reporter^84^ or the *her6:*mCherry reporter^85^, both of which are expressed in the CS after ∼36 hpf. We detected a close physical interaction between *sox10:*DsRed*+* cells and the CS as early as 50 hpf (Fig. 6C), after the glands had fully formed. The closest cells appear to turn off *sox10* shortly thereafter, as they became harder to find, though lineage-traced crest were present in the vicinity of the CS up to 7 dpf (Fig. 6D-E). Strikingly, *sox10* mutants lack *SOX10*:Cre lineage-labeled cells around the CS at all stages examined (Fig. 6D-E). This is consistent with the complete or near-complete loss of many neural crest cell sublineages previously reported in *sox10* mutant models^6,7,86^. Mutant VenusPEST+ CS cells are less organized, and mutant gland volume is larger (p<0.0001 at 58 and 72 hpf, p=0.003 at 96 hpf, ns at 168 hpf; unpaired t-tests; Fig. 5E’). These patterns suggest that *sox10+* crest-derived cells may act locally to restrain CS growth and *stc1a* expression to regulate embryonic calcium homeostasis.

### *stc1a* is epistatic to *sox10* and the proximate cause of the mineralization deficit

Our results thus far suggested that the absence of *sox10*+ cells leads to unrestrained growth and Stc1a production by the Corpuscles, in turn inhibiting NaR cell proliferation and preventing sufficient calcium uptake for mineralization. To test this model, we performed an epistasis assay of *stc1a* on the *sox10* mutant background using the previously reported *stc1a^mi6^*^10^ mutant^42^. *sox10^ci30^*^20^*; stc1a^mi610^* double mutants present both the trademark lack of pigmentation and underdeveloped inner ears of *sox10* single mutants alongside the characteristic cardiac edema of *stc1a* mutants (Fig. 7A), supporting that these phenotypes are genetically independent. However, bone mineralization was strikingly improved in double mutants relative to *sox10* single mutants at 4 dpf (Fig. 7B). Eighty percent of the double mutants (24 out of 30) stained with Alizarin red: 13 weakly, 10 intermediate, and 1 strongly (Fig. 7E, also see Fig. S5B for examples). It is worth noting that the presence of cardiac edema in the double mutants may have compromised bone formation in some individuals. For comparison, among 23 *sox10^-/-^; stc1a^+/+^* individuals, 14 had no staining, 5 had weak staining, 3 intermediate, and 1 strong (Fig. S5B; p=0.0206, Chi-square test). In the original *sox10^ci3020^* single mutant crosses, only 3/48 single mutants showed intermediate or weak staining; the other 45 had none (Fig. S5A), suggesting the presence of genetic modifiers. We further noted significant improvement in NaR cell number and calcium content in the double *sox10; stc1a* mutants relative to *sox10* single mutants (Fig. 7C’-D), further supporting that *stc1a* is epistatic to *sox10* in mineral regulation.

**Figure 7.**
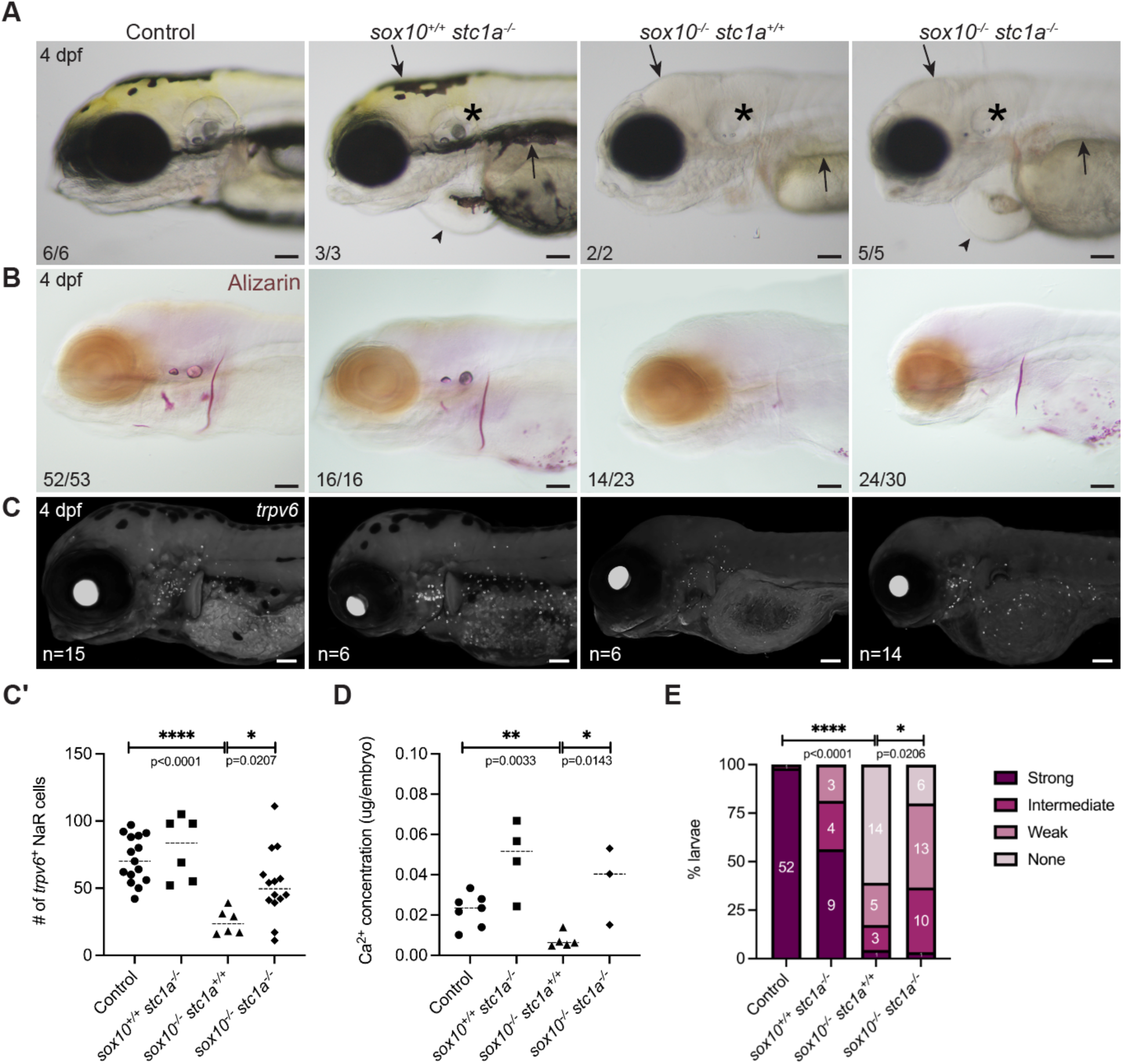
*stc1a* is epistatic to *sox10* in control of systemic calcium content. **(A)** Brightfield images of *sox10* and *stc1a* controls and mutants at 4 dpf. Double mutants phenocopy the loss of pigment (arrow) and the inner ear malformations (asterisk) of single *sox10* mutants and the cardiac edema of the *stc1a* mutant (arrowhead). **(B-C’)** Loss of *stc1a* on the *sox10* mutant background improves mineralization **(B)** and the number of *trpv6+* ionocytes **(C)** at 4 dpf, quantified in C’ (unpaired t-test; p=0.0207, df=18). Dashed bars indicate the median. **(D)** Calcium quantification shows an increase (unpaired t-test; p=0.143, df=6) in calcium levels in *sox10^-/-^; stc1a^-/-^* compared to *sox10^-/-^.* Dashed bars indicate the median. **(E)** Quantitation of mineralization levels in *sox10; stc1a* clutches grouped based on the intensity of the Alizarin red staining. There was a significant increase in the proportion of double mutants with detectable mineralization compared with *sox10* single mutants (Chi-square; p=0.0206, df=3). In C’-E, ‘control’ includes wild-type and heterozygous larvae. Scale bars: 100 µm.

## Discussion

### Novel Sox10 requirement in bone mineralization

This study challenges the decades-old paradigm that Sox10 is not required for skeletal development by revealing a previously undescribed, indirect role in mineralization. Two independent *sox10* mutant lines exhibit delayed and reduced mineralization of all bones, no matter their embryonic origin or ossification type. Mutant osteoblasts appear to differentiate normally (Fig. 2D) and gradually lay down ECM to create typically-sized bone templates (Fig. 2C-E). However, their transcriptomes may be subtly altered: we detected changes in whole-body mRNA levels of genes encoding osteoblast-enriched enzymes involved in regulation of phosphate availability and homeostasis (*phospho1*, *alpl*, *enpp1*, *entpd5*) or bone accessory proteins (*spp1* (osteopontin) (Fig. S2A-B). We posit that these shifts may reflect secondary transcriptional responses of osteoblasts to the major systemic calcium deficit or to changes in inorganic phosphate availability incurred by the lack of calcium. Reduced levels of some of these factors may exacerbate the mineralization defect in *sox10* mutants, as other studies have demonstrated that partial or complete genetic loss of some of these accessory proteins and enzymes can lead to decreased bone mineral density and/or mineralization deficits^62–65^.

We noted with interest the changes in phosphate regulators, given that we did not measure any consistent differences in mutants’ total phosphate content by a colorimetric assay (Fig. S3B). It is possible that the assay is insufficiently sensitive or overwhelmed by maternally deposited yolk stores^26–28^. However, how osteoblast-engendered inorganic phosphate intended for hydroxyapatite formation is managed when calcium is not available is an intriguing question. Of note, in our comparison of bone stains, we observed recovery of Von Kossa staining before that of Alizarin red, Calcein, or OsteoImage (hydroxyapatite). In Von Kossa staining, silver cations from the silver nitrate staining solution interact with calcium phosphate to produce a yellowish silver phosphate, which subsequently blackens surrounding organic matter^52,87,88^. It is possible that the early recovery of this stain reflects a reaction with inorganic phosphate accumulating in the bone matrix due to the calcium deficit.

How calcium uptake and bone mineralization begin to recover in *sox10* mutants is still an open question. One possibility is that other endocrine hormones come ‘online’ and begin to counteract elevated Stc1a activity. Parathyroid hormone and vitamin D are reported to have hypercalcemic properties in fish as well as in mammals, acting to increase Trpv6-mediated calcium uptake^21,89,90^. Zebrafish lack parathyroid glands, but express parathyroid hormones in the central nervous system and sensory neuromasts^91^. Similarly, fish synthesize vitamin D as early as 3 dpf in response to decreased environmental calcium^89^. Other physiological changes are occurring in fish larvae at the same time that mineralization begins to recover, including maturation of the digestive tract and auxiliary endodermal organs^92^ and depletion of the yolk^26^. Though *sox10* mutants lack an enteric nervous system^7^ and are not fed in our experiments, it is possible that passage of embryo medium through the digestive tract allows calcium uptake through intestinal enterocytes, contributing to the mutants’ partial recovery. We have also observed ectopic calcium/hydroxyapatite deposits in the yolk area of mutants at 3 and 4 dpf that begin to resolve coincident with the onset of bone mineralization (Fig. 1C-D). The calcium in those deposits could conceivably be remobilized and made available for forming bones as the yolk is depleted. Two other zebrafish mutants that lack mineralization during larval stages (*msp*^70^, *her9*^58^) also naturally recover to some extent, supporting robustness or complementarity in mechanisms driving calcium uptake for skeletal mineralization.

### Sox10 drives bone mineralization indirectly through interactions with endocrine glands involved in calcium homeostasis

The most striking finding from the whole-body transcriptional analysis was the tripled *stc1a* mRNA levels in *sox10* mutants (Fig. 5A). High Stc1a blocks proliferation of *trpv6+* ionocytes^42^, reducing calcium uptake. That elevated *stc1a* is the major driver of the calcium deficit in *sox10* mutants was confirmed by our epistasis studies (Fig. 7). However, whether the increase in *stc1a* mRNA is attributable solely to the higher cell number in the mutant Corpuscles (Fig. 5C-D) or also to a percell increase in transcription is not yet clear. Previous studies have shown that high external calcium upregulates *stc1a* transcription at least in part via the Calcium-Sensing Receptor (CaSR), which is also expressed in the CS^77^. Aberrant activity of CaSR in the absence of *sox10*+ lineage cells could therefore potentially boost *stc1a* transcription. In support of the idea that the *stc1a* increase is more complex than just increased CS cell number, another mineral-regulating hormone enriched in the CS, *fgf23*^93,94^, is downregulated in *sox10* mutants (Fig. S2A) despite the increased size of the Corpuscles. Fgf23 has anti-hypercalcemic effects similar to Stc1a, reducing Ca^2+^ uptake in conditions of high systemic calcium, in addition to regulating phosphate homeostasis^77,95,96^; its low expression in *sox10* mutants is consistent with their calcium deficit^77^. Published scRNAseq data^74^ shows that Corpuscle cells also express receptors for other endocrine factors involved in mineralization between 2 and 4 dpf, including receptors for calcitonin (*calcr*), cortisol (*nr3c1*), vitamin D (*vdrb)*, Fgf23 (*fgfr1b*), and Msp (*mst1rb*). It remains to be seen how these pathways are affected in the absence of *sox10*+ cells and whether they are involved in *stc1a* upregulation.

Why the Corpuscles, derived from a *sox10*-negative mesodermal lineage, are so profoundly affected by loss of *sox10* is not fully resolved. We did not observe an increase in *stc1a*+ cell number before 2 dpf, i.e., only after the glands had fully extruded from the pronephros, ruling out expanded CS specification as the explanation for the larger glands (as previously found in other mutant lines^80,97^). Our experiments instead revealed that a *sox10*+ sublineage interacts with these glands post-extrusion, and that these NC-derived cells are missing in *sox10* mutants (Fig. 6C-E), like so many other crest derivatives^6–9^. A tantalizing possibility is that they may be precursors of the sympathetic neurons that will innervate the CS in adults^79,98^. Sympathetic neurons derive from *sox10*+ neural crest, in particular from Schwann cell precursors (SCPs)^99^. Differentiating Schwann cells and SCPs are thought to be the predominant *sox10*+ cell types lining the trunk sensory and motor axon tracts that pass by the Corpuscles^3,99^, from which we see cells emerging to contact the glands directly (Fig. 6C). Schwann cells and SCPs are largely absent in *sox10* mutant fish and mice^7,10^. Interestingly, hallmark signs of sympathetic neuronal differentiation in the trunk are not evident in wild-type zebrafish until around 7 dpf^100^, well after this CS phenotype arises. The regulatory interaction between the *sox10*+ lineage cells and the CS is thus expected to be non-neuronal in nature at these early stages. Though mutant lethality makes it challenging to study the onset of sympathetic control, we expect that the requirement for *sox10*+ lineage cells in managing stanniocalcin production and/or secretion and thus calcium homeostasis persists throughout the lifespan.

Humans and other mammals do make stanniocalcin hormones, but we do not develop a gland homologous to the Corpuscles of Stannius^35^. If loss of *sox10* in fish impacts mineralization solely through dysregulation of Corpuscle development and function, as our data support, it is conceivable that mammals lacking *Sox10* will show no equivalent signs of mineral dysregulation. However, our studies also prompt the more general notion that crest-derived cells destined to become part of the sympathetic nervous system may make contact with and begin regulating their target organs’ growth and activity earlier in embryonic development than previously appreciated. This could potentially drive physiological and endocrinological symptoms in individuals with congenital neurocristopathies caused by deficient crest production or survival^101^.

## Materials & Methods

### Zebrafish husbandry and lines

Zebrafish embryos were grown at 28.5°C in standard embryo medium (EM) ^102^ unless otherwise noted: 15 mM NaCl, 0.5 mM KCl, 1 mM CaCl_2_•2H_2_O, 0.15 mM KH_2_PO_4_, 0.06 mM NaH_2_PO_4_ and 1 mM MgSO_4_•7H_2_O. Published mutant and transgenic lines used here include *sox10^ci3020^* ^49^ *sox10^m618^* ^51^, *stc1l^mi610^* ^42^, *Tg(Hsa.RUNX2:mCherry*)*^zf3244^* (alias *RUNX2:mCherry*)^59^, *Tg(sp7*:*EGFP*)*^b121260^*, *Tg(Ola.Bglap:EGFP*)*^hu4008^*(alias *osc:EGFP*) ^29^, *Tg(Mmu.Sox10-Mmu-Fos:Cre)^zf384^* (alias *SOX10*:*Cre*) ^48^, *Tg(EPV.TP1-Mmu.Hbb:Venus-Mmu.Odc1)^s940^*(alias *Tp1:VenusPEST*) ^84^, *Tg(fli1:EGFP)^y1^*^103^, *Tg(Xla.Eef1a1:loxP-DsRed2-loxP-EGFP*)*^zf284^*(alias *ef1a:DsRed>EGFP*) ^48^, *Tg(actb2:LOXP-BFP-LOXP-DsRed)^sd27^* (alias *actb2:*BFP*>*DsRed) ^82^ and *Tg(her6:mCherry)^sd64^* ^85^. Lines were maintained as hetero- or hemizygotes.

### Bone staining

For all fixed bone stains, zebrafish larvae were fully anesthetized with MS-222 (aka Tricaine, Syndel, USA) at the desired stage and then fixed in 2% paraformaldehyde (PFA) (250 ml embryo medium, 250 ml 4% PFA, and 500 ml PBS with 0.1% Tween) overnight at 4°C or for 1 hour at room temperature. For Alizarin red-only staining, following fixation, larvae were rinsed twice in 25% glycerol in 0.5% KOH for 10 minutes each and stained with 0.01% Alizarin in 25% glycerol/100 mM Tris pH 7.5 for 4 hours at room temperature. They were then bleached for 10 minutes in 3% H_2_O_2_ in 0.5% KOH under a light source. Specimens were stored and imaged in 50% glycerol in 0.5% KOH or 100% glycerol immediately to prevent fading (adapted from^104^). Combined Alcian blue and Alizarin red staining was performed as described previously^105^. For Von Kossa staining, fixed embryos were rinsed with deionized water and stained with 2.5% silver nitrate solution (Abcam ab150687) under a light source for 20 minutes. The reaction was stopped with 5% sodium thiosulfate to prevent overstaining, and larvae were imaged immediately^106,107^. For the Osteoimage^TM^ Mineralization Assay (Lonza PA-1503), we followed the manufacturer’s protocol after fixing. Briefly, fixed larvae were rinsed with diluted wash buffer then stained in diluted Staining Reagent for 30 minutes at room temperature in the dark. Before imaging, they were rinsed three times with wash buffer for five minutes each. For live staining, larvae were incubated in Alizarin red (0.03 mg/ml in 30 ml EM) for 2 hours at 28.5°C or in Calcein green (0.1 mg/ml in 30 ml EM) at 28.5°C overnight^55^. For each round of each bone staining experiment, a minimum of six individuals were stained and imaged per genotype/stage/group.

### Calcium and phosphate supplementation and depletion treatments

For calcium treatments, the amount of CaCl_2_•2H_2_O was increased two or ten-fold for 2 mM and 10 mM treatments, respectively, completely removed (0 mM), or decreased to 0.02 mM^70^). For the high phosphate treatment (adapted from ^62^), the concentrations of KH_2_PO_4_ and NaH_2_PO_4_ were raised to 0.5 mM and 9.5 mM, respectively, to increase the total PO_4_^3-^ to 10 mM, therefore maintaining the proportional K^+^/Na^+^ ratio as in the control EM. The ‘No PO_4_^3-^’ treatment included neither KH_2_PO_4_ nor NaH_2_PO_4_ in the media. A minimum of six control and six mutant larvae were used per treatment group, and all treatments were repeated at least twice.

### Whole mount in situ hybridization and immunostaining

cDNAs for *stc1a*, *trpv6*, *igfbp5a, col10a1a*, *phospho1, sparc, spp1, runx2a, and runx2b*^108^ were amplified by Herculase II Fusion DNA Polymerase (Agilent) (see Table S1 for primer sequences) and inserted into the pCR-Blunt II-TOPO vector (ThermoFisher). After sequence confirmation and linearization by restriction digest, antisense probes were synthesized from each plasmid using Sp6 or T7 polymerase and digoxigenin (DIG)-tagged nucleotides (Roche). Colorimetric and fluorescent *in situ* hybridizations were performed as described previously^109^. Colorimetric *in situs* were developed with either NBT-BCIP or BM Purple (Sigma-Aldrich), whereas fluorescent *in situs* were developed with TSA Cyanine 3 (Akoya Biosciences). Immunostaining was performed as described^49^. Primary antibodies included anti-Na^+^/K^+^ ATPase (1:400, DSHB α5) and anti-Sox10 (1:500, Genetex GTX128374), used with AlexaFluor 647-conjugated goat anti-mouse and donkey anti-rabbit secondary antibodies (1:250, ThermoFisher A32728 and B40956). In both procedures, permeabilization steps were skipped for markers limited to surface expression (*trpv6, igfb5a* and α5). A minimum of six control and six mutant larvae were stained and imaged for each marker, and the experiments were repeated at least twice.

### Semi-quantitative reverse-transcriptase PCR (rt-PCR)

rt-PCRs were performed to estimate transcript levels of mineralization-associated genes in *sox10^ci^*^302^*^0^* mutants. Each sample consisted of 10-15 mutant and 10-15 stage-matched wild-type controls that were pooled at 4 dpf and frozen at −80°C. RNA was extracted using the RNAqueous-4PCR Total RNA Isolation Kit (Invitrogen), and equivalent amounts were used to synthesize cDNA with the High-Capacity cDNA Reverse Transcription Kit (Applied Biosystems). rt-PCR was run with a minimum of three biological replicates per genotype, and *eef1g* expression was used for normalization (following ^49^). Band intensity was quantified with Image Lab (BioRad) and analyzed with Prism 10 (GraphPad). Primers, product sizes, and cycling conditions for each gene are listed in Table S2.

### Quantification of mineral content

Whole-body Ca^2+^ and PO_4_^3-^ contents were quantified using colorimetric assay kits (Abcam ab102505 and ab65622). For Ca^2+^ measurements, 10-15 larvae were pooled at the desired stage in an Eppendorf tube without any liquid, dehydrated at 60°C for 1 hour, then digested for at least 4 hours in 125 μL of freshly prepared 1M HCl in an Eppendorf Thermomixer set at 95°C and 750 rpm. The samples were then centrifuged at 4°C for 45 minutes at 15,000 rpm. Supernatants were distributed on a clear 96-well polystyrene flat-bottomed plate alongside the standard curve reagents prepared according to the protocol provided with the kits. Absorbances were measured on a SpectraMax M5 plate reader. The same procedure was followed for PO_4_^3-^ quantification, with the modification that the supernatants were diluted in deionized water to avoid precipitation.

### Imaging and image analysis

Skeletal stains, brightfield images and colorimetric *in situs* were imaged on a Zeiss SteREO Discovery.V8 or Zeiss Axioimager.Z1 microscopes, whereas fluorescent *in situs*, fluorescent bone stains, live transgenic fish, and immunostained specimens were imaged on a Nikon C2 confocal. *trpv6+/igfbp5a+* ionocytes and *stc1a+* cells were quantified using the ‘spots’ option in Imaris 10.1.1. CS volumes were measured with the surface labeling option in Imaris 10.1.1. A minimum of six replicates were counted for each genotype/stage combination.

### Data analysis

Data analysis was performed with GraphPad Prism (Version 10.2.3). p-values were calculated with Chi-square tests or unpaired two-tailed t-tests as noted in the figure legends.

## Acknowledgments

We are grateful to members of the Barske lab for helping with molecular biology experiments and imaging; Kristina Preusse and Benjamin Liou for assistance with mineral quantification; Evan Brooks and Samantha Brugmann for helping to set up Von Kossa and OsteoImage staining; Colin Kenny, Chunyue Yin, Claire Arrata, and Gage Crump for sharing fish lines; Flynn Littleton, Eric Alley and the CCHMC Division of Veterinary Services for fish care; and Josh Gross, James Nichols, Jessica Nelson, and Rolf Stottmann for helpful discussions and/or manuscript suggestions. Funding for this project was provided to L.B. by Cincinnati Children’s Center for Pediatric Genomics and the Cincinnati Children’s Research Foundation; funding for L.S. and K.C.S. was provided by BBSRC grant BB/S015906/1 to R.N.K.

## Author Contributions

The project was conceived by S.G. and L.B. Zebrafish experiments were performed by S.G., S.P., S.M. L.S., K.C.S., and L.B. Crucial fish lines and guidance were provided by C.D. and R.K. Writing and interpretation were performed primarily by S.G. and L.B. with input from C.D. and R.K.

