## Supplemental Data for "Sox10 is required for systemic initiation of bone mineralization"

### Supplementary Data:

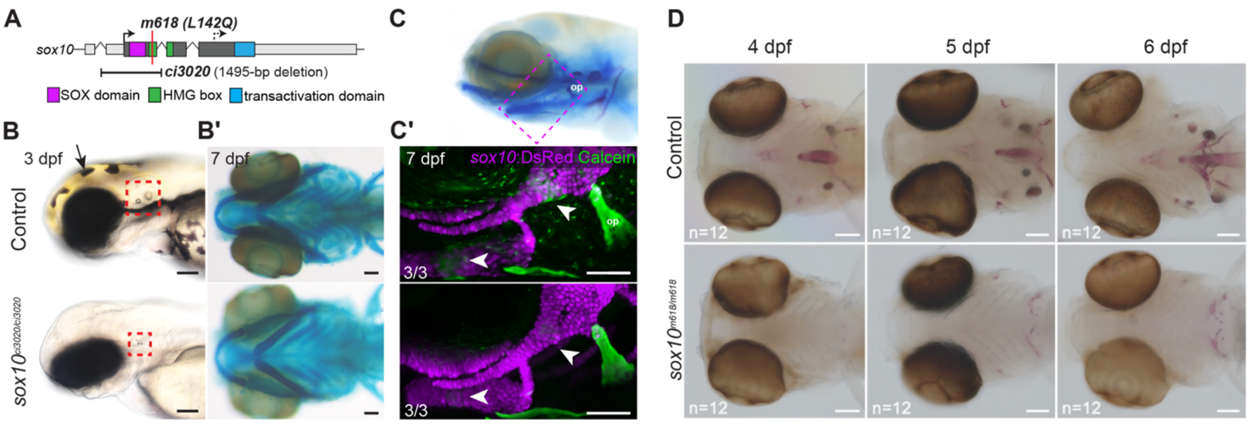


**Figure S1. Validation of *sox10* null allele. (A)** Schematic representation of the *ci3020* and *m618* mutations in the *sox10* gene body. Dotted arrow indicates the next in-frame methionine downstream of the *ci3020* deletion. **(B-B')** $sox10^{ci3020/ci3020}$ mutants lack melanocytes (arrow) and yellow xanthophores and have inner ear and otolith malformations (dashed red box) (B) but form a normal cartilaginous craniofacial skeleton (Alcian blue staining) (B'). Scale bars: 100 µm. **(C-C')** Lateral view of control embryo stained with Alcian blue and Alizarin red, showing the position of image in C'. $sox10^{ci3020/ci3020}$ mutants still show deficient endochondral mineralization at 7 dpf, while intramembranous bones (e.g. op) have begun to catch up. Arrowheads indicate sites of bone collars on the hyomandibular and ceratohyal cartilages. *sox10*:DsRed (magenta) marks chondrocytes, and Calcein green labels mineralized bone. Scale bar: 100 µm. **(D)** The missense $sox10^{m618/m618}$ mutant also shows deficient mineralization at 4-6 dpf by Alizarin red staining. Scale bar: 100 µm.

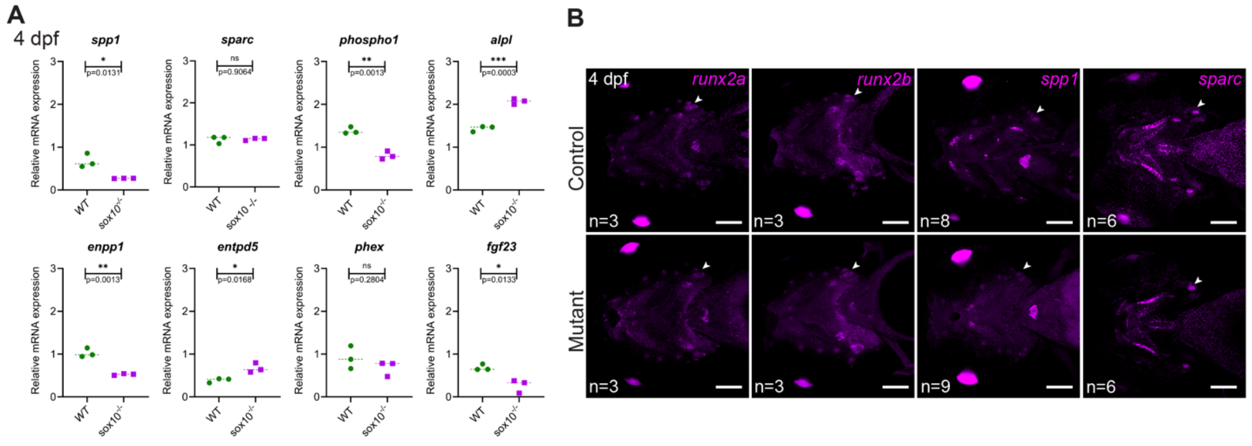


**Figure S2. Altered expression of genes encoding mineralization enzymes and matrix proteins in *sox10* mutants. (A)** Semi-quantitative rt-PCRs were performed on cDNAs made from three pools of 10-15 wild-type (WT) or mutant embryos collected at 4 dpf. The analysis revealed significant increases in *alpl* and *entpd5* (unpaired t-tests: p=0.0003, and p=0.017, respectively, df=4 for both), decreases in *spp1*, *phospho1*, *enpp1*, and *fgf23* (unpaired t-tests: p=0.013, 0.001, 0.001, and 0.013, respectively, df=4 for all), and no changes in *sparc* or *phex* (unpaired t-tests: p=0.906 and p=0.284, respectively, df=4 for both). **(B)** Fluorescent *in situ* hybridizations show no changes in *runx2a* or *runx2b* but reduced *spp1* and *sparc* expression. These phenotypes were consistent across all samples imaged. Arrowheads pointing at op. Scale bars: 100 μm.

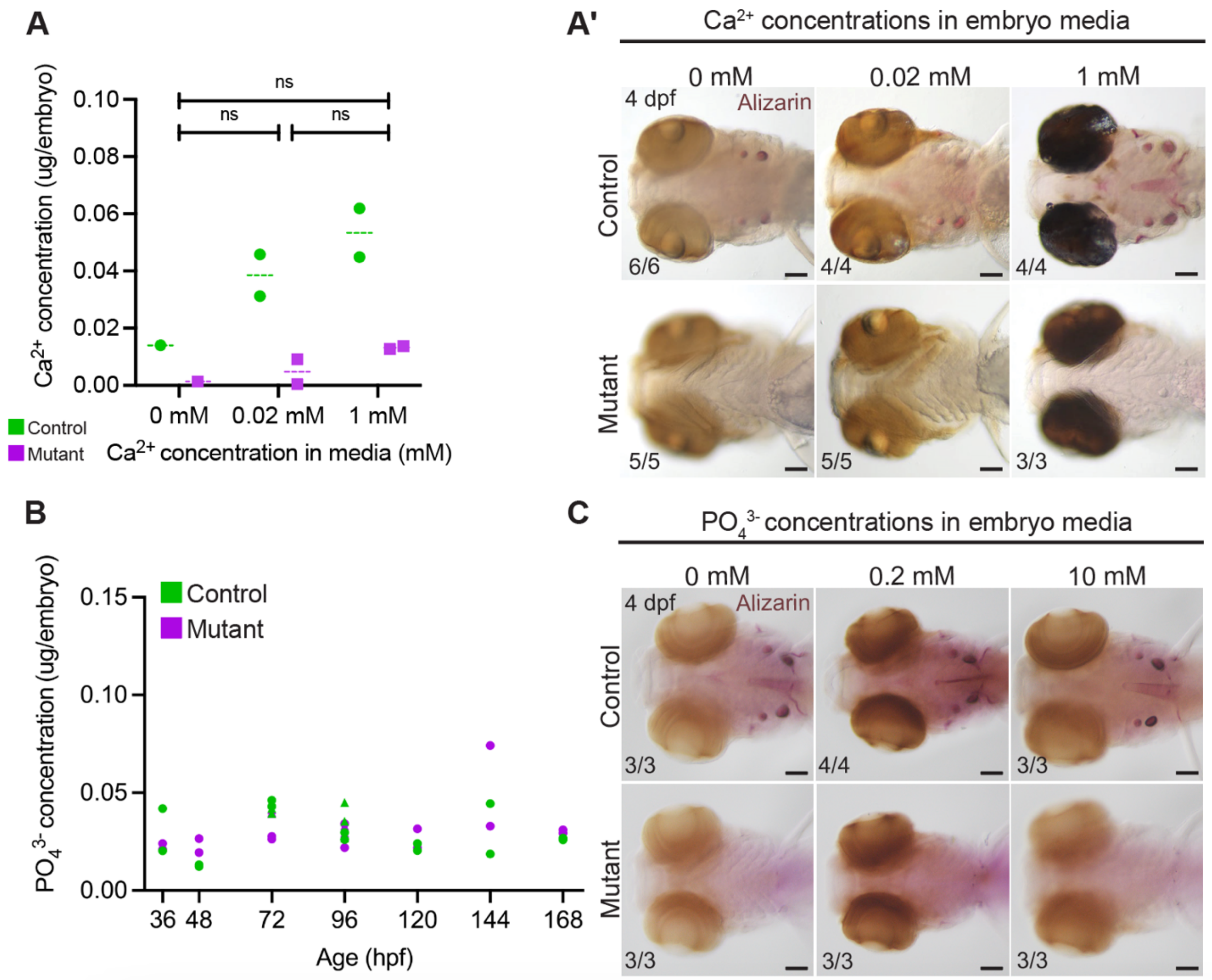


**Figure S3. Neither low calcium nor high or low phosphate improve bone mineralization in *sox10* mutants. (A)** Reduction or complete removal of calcium from the embryo media impairs mineralization in control larvae and does not rescue the mutant phenotype (unpaired t-tests: 0.02 vs. 0 mM: p=0.519, df=2; 1 vs. 0 mM: p=0.347, df=2; 1 vs. 0.02 mM: p=0.700, df=2). Otoliths, made from calcium carbonate rather than hydroxyapatite, still form in controls in the absence of external calcium. Calcium levels of mutants and controls raised to 4 dpf in different media are quantified in A', matching the corresponding Alizarin staining patterns. **(B)** Phosphate levels do not overtly differ between controls and *sox10* mutants throughout larval development (1.5-7 dpf) (unpaired t-tests: 36 hpf: p=0.516, df=2; 48 hpf: p=0.104, df=2; 72 hpf: p=0.046, df=8; 96 hpf: p=0.550, df=8; 120 hpf: p=0.474, df=2; 144 hpf: p=0.462, df=2; 168 hpf: p=0.005, df=4). **(C)**

Increasing (10 mM) or removing (0 mM) phosphate from the embryo media does not improve mineralization in *sox10* mutants. Scale bars: 100 µm.

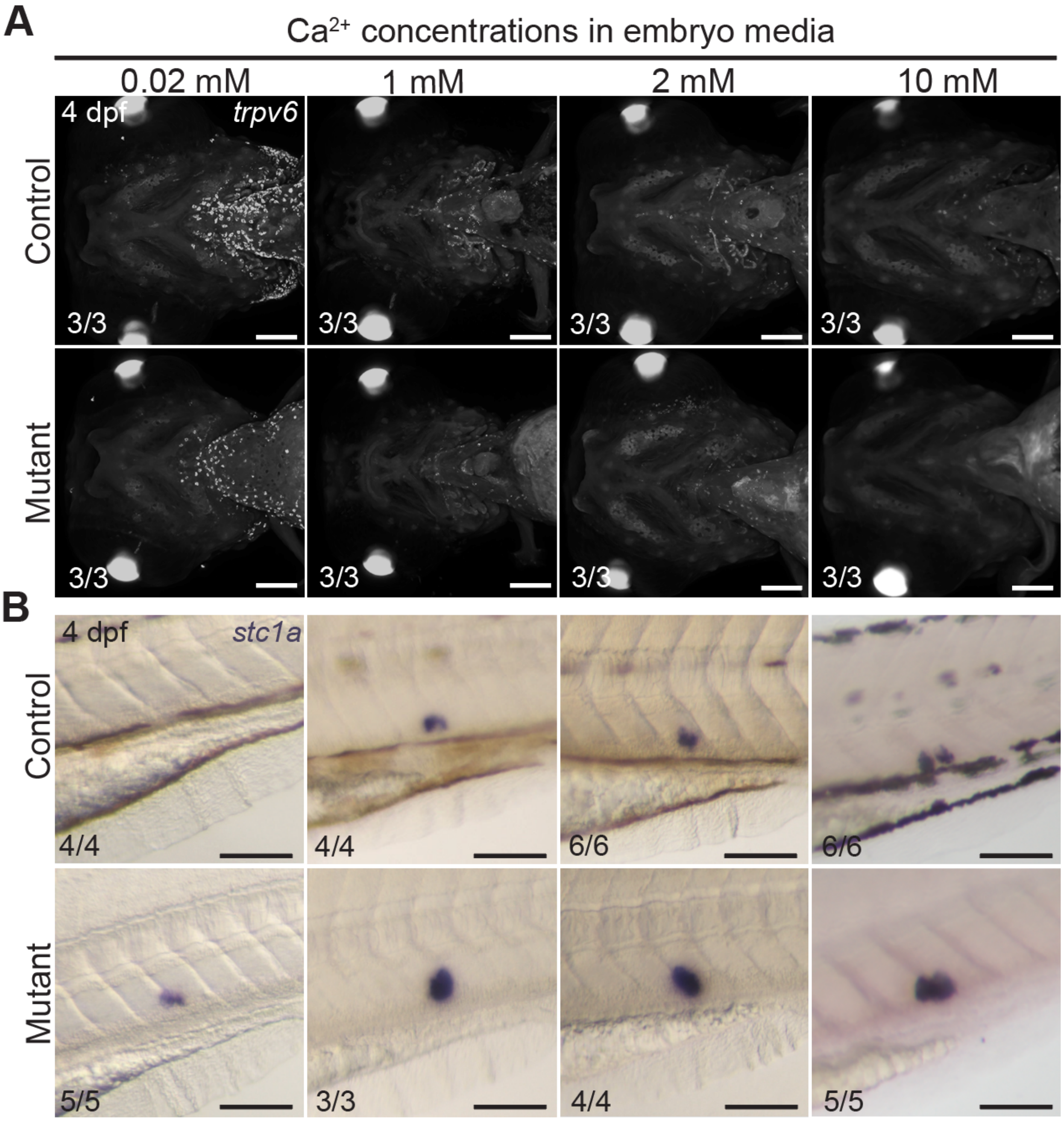


**Figure S4. *sox10* mutants still sense and respond to ambient calcium levels. (A)** Fluorescent *in situ* hybridization confirms clear increases and decreases in the number of *trpv6*+ cells in both control and mutant larvae raised to 4 dpf in high (2 or 10 mM) or low (0.02 mM) environmental calcium, respectively, compared to standard embryo media (1 mM). **(B)** *stc1a* mRNA levels are reduced in both control and mutant larvae raised to 4 dpf in low environmental $Ca^{2+}$ compared to those kept in standard medium, though transcription is not shut off completely in mutants. Levels appear marginally increased in controls raised in high $Ca^{2+}$ but are not overtly changed in mutants. Scale bars: 100 µm.

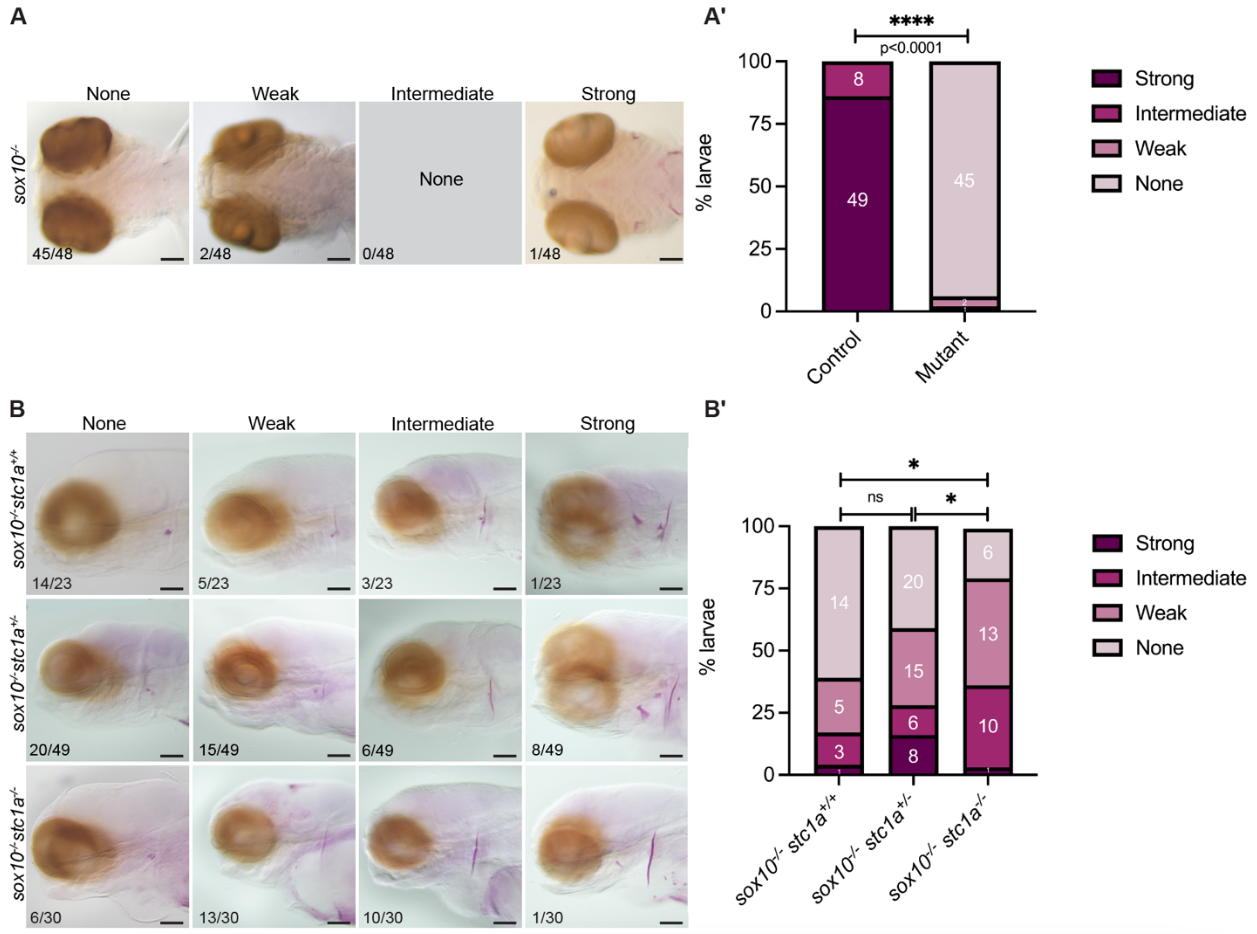


**Figure S5. Spectrum of mineralization intensities in *sox10* mutants at 4 dpf. (A-A')** By Alizarin red staining, 92% (45 of 48) of *sox10* mutants imaged at 4 dpf showed a complete lack of mineralization. The remaining 8% had either faint (classified as 'weak'; 2 of 40) or strong (1 of 40) Alizarin staining. All controls were classified as intermediate or strong. There is significantly less mineralization in the mutants compared to controls (Chi-square; p<0.0001, df=3). **(B-B')** Partial genetic reduction of *stc1a* improved mineralization in *sox10* mutants, but a more significant improvement (Chi-square; p=0.0206, df=3) was observed upon complete loss of *stc1a*, as 83% of the double mutant larvae (24 out of 30) had some mineralization. The incidence of faint Alizarin staining in $sox10^{-/-}$; $stc1a^{+/+}$ larvae produced by incrossing double heterozygotes was higher than in the single *sox10* mutants quantified in A, possibly due to genetic background effects. Classes: 'strong': well-stained opercle, cleithrum and teeth; 'intermediate': well-stained cleithrum, no opercle or teeth; ‘weak’: faint cleithrum, no opercle or teeth; ‘none’: no Alizarin red staining. Scale bars: 100 μm.

**Table S1: *In situ* probe primers**

| Genes | Primers (5'→ 3') | Linearization Enzyme | RNA polymerase |
|---|---|---|---|
| *trpv6* | GGCAACGACCACACCATAAA | EcoRV | Sp6 |
| | TGGGCCATTATGTCATTGCG | | |
| *col10a1a* | ACCAGCCTTACTCCGTGAAA | EcoRV | Sp6 |
| | GGCTCACCTTTCTGACCAGT | | |
| *stc1a* | ATGCTCCTGAAAAGCGGATTTCTT | EcoRV | Sp6 |
| | AGGACTTCCCACGATGGAGCGTTT | | |
| *igfbp5a* | AGTTTGTATGCTCTGGTGGC | EcoRV | Sp6 |
| | CCATTAAAGTCAGTGCCCGG | | |
| *phospho1* | CCGCTTCCTGATGTTCTTCG | EcoRV | Sp6 |
| | GTCCTCACCACTCTTCCAGG | | |
| *sparc* | CTTCTTCCTGTTCTGCCTCG | EcoRV | Sp6 |
| | ATAAGAGGAGCACGCAGAGG | | |
| *spp1* | CCACGCCAACAGAATCGAAT | EcoRV | Sp6 |
| | TCTCCTGGCTTTCTGTGCTT | | |
| *runx2a* | CGCGTGTGTTTGTTTGTTCC | EcoRV | Sp6 |
| | CATGGTGGTCTGGCGTAAAC | | |
| *runx2b* | CGGTGAAGATGAACGACGTG | XbaI | T7 |
| | GCCAGGGAAAGGACTCAAGT | | |

**Table S2: rt-PCR primers**

| Genes | Primers (5’à 3’) | Product size (bp) | Conditions* |
|---|---|---|---|
| *stc1a* | CACGGTTCTCATCCAACACC<br>GCGCTTAATGGTCTGGAACA | 245 | 15s/30s/30s 56°C 30x |
| *spp1* | CAGCAAGCAGTTCAGAGAGC<br>CTGCCTCCTCAGTGTCATCT | 162 | 15s/30s/30s 56°C 30x |
| *sparc* | TCCAACTGAAGAGGAGCCAG<br>ATAAGAGGAGCACGCAGAGG | 164 | 15s/30s/30s 56°C 30x |
| *enpp1* | GTGCTTCTTCTTGGCTTGCT<br>CAGGTGCCTTCATTTACGCA | 167 | 15s/30s/30s 56°C 28x |
| *entpd5* | CGCTACCGTGCAATCAATCA<br>TGGAAGCTCAACTGGGTCTT | 195 | 15s/30s/30s 56°C 28x |
| *phex* | CTCTGTCTAGGCCCACACTG<br>TATCTCCACTTCTCACGGGC | 226 | 15s/30s/30s 56°C 28x |
| *phospho1* | TTGCCCGTCTCTTACACTGT<br>GTCCTCACCACTCTTCCAGG | 142 | 15s/30s/30s 56°C 30x |
| *fgf23* | ACATCCACCTTCAGACCACT<br>TCCTCCTGTATCCATGCACA | 185 | 15s/30s/30s 56°C 30x |
| *alpl* | AGTTTCCAGAGCAAGAGAAGC<br>TCTGTCCACTCAACTGACCC | 189 | 15s/30s/30s 56°C 30x |
| *trpv6* | CAGGCATGTACTTCCGCAAA<br>ATCCTGAGCCAGCAAGAAGT | 227 | 15s/30s/30s 58°C 35x |
| *eef1g* | TCGTCTGAAGATTGCGAGTG<br>ACCCTGGTAAGCTGGAACCT | 109 | 15s/30s/30s 56°C 25x |

*Format: 15s denaturation, 30s annealing, 30s elongation, annealing temperature, # cycles
